# Deep learning with feature embedding for compound-protein interaction prediction

**DOI:** 10.1101/086033

**Authors:** Fangping Wan, Jianyang (Michael) Zeng

## Abstract

Accurately identifying compound-protein interactions *in silico* can deepen our understanding of the mechanisms of drug action and significantly facilitate the drug discovery and development process. Traditional similarity-based computational models for compound-protein interaction prediction rarely exploit the latent features from current available large-scale unlabelled compound and protein data, and often limit their usage on relatively small-scale datasets. We propose a new scheme that combines feature embedding (a technique of representation learning) with deep learning for predicting compound-protein interactions. Our method automatically learns the low-dimensional implicit but expressive features for compounds and proteins from the massive amount of unlabelled data. Combining effective feature embedding with powerful deep learning techniques, our method provides a general computational pipeline for accurate compound-protein interaction prediction, even when the interaction knowledge of compounds and proteins is entirely unknown. Evaluations on current large-scale databases of the measured compound-protein affinities, such as ChEMBL and BindingDB, as well as known drug-target interactions from DrugBank have demonstrated the superior prediction performance of our method, and suggested that it can offer a useful tool for drug development and drug repositioning.

## 1 Introduction

Identifying compound-protein (or drug-target) interactions is crucial in drug discovery and development in that it provides valuable insights into understanding efficacy and off-target adverse side-effects of drugs [1] [2]. Also, inspired by drug polypharmacology, i.e., single drugs can interact with multiple targets [3], drug developers are seeking to better characterize compound-protein interactions to find new uses of existing drugs (also called drug repositioning or drug repurposing) [3] [4] [5], as it can significantly reduce the time and cost required in drug development [6].

Numerous computational methods have been proposed to predict potential compound-protein interactions *in silico* to narrow the large search space of possible interacting compound-protein pairs and facilitate the drug discovery and development process [7] [8] [9] [10] [11] [12] [13] [14]. Although a number of successful predictions have been achieved, there is still much room for improvement in existing computational prediction approaches. First, most of the previous prediction methods only use a simple and direct representation of features from the labeled data (e.g., known compound-protein interactions and available structural information of proteins) to measure the similarities between compounds or proteins, and infer the unknown compound-protein interactions. For instance, a drug-protein interaction profile kernel [11] and a graph-based method SIMCOMP [15] were used to compare two drugs or compounds, and the normalized Smith-Waterman score [13] was typically applied to assess the closeness between two targets (proteins). On the other hand, the huge amount of available unlabelled data of compounds and proteins can also provide an implicit and useful representation of features that can be effectively used to define their similarities. Such an implicit representation of protein or compound features encoded by the large-scale unlabelled data has not been well exploited in most of existing methods to predict new compound-protein interactions. Second, an increasing number of deposited drug-target interactions or compound-protein binding affinities (e.g., one million bioassays over two million tested compounds and ten thousand protein targets in PubChem [16]) raises serious scalability challenge to previous prediction methods. For example, many similarity-based methods [13] [9] require the computation of pair-wise similarity scores between proteins, which is generally impractical in the setting of large-scale data. Third, most of existing prediction methods have difficulty in predicting new drug-target pairs (i.e., both drug and target in the pair have no known interaction knowledge). The aforementioned computational challenges generally require one to develop a more effective scheme to accurately capture the hidden features of proteins and compounds and a more advanced learning model to accomplish the prediction task.

In the machine learning community, representation learning and deep learning are now two popular techniques for efficient feature extraction and addressing the scalability problem in large-scale data analysis. Representation learning aims to automatically learn data representations (also called features) from relatively raw data that can be effectively exploited by downstream machine learning models to improve the task performances [17] [18] [19]. Representation learning for proteins and compounds have been previously explored in [20] [21] to achieved promising results in other prediction tasks. Deep learning, normally with fast GPU implementation, which extracts high-level abstractions of features from input data by typically using several layers of non-linear transformations, has become one of the dominant methods in numerous complex tasks with large-scale data in many data science fields, such as computer vision, speech recognition, natural language processing and game playing [22] [23] [24] [25] [26]. In the literature, several deep learning models have been proposed for compound-protein binding prediction [27] [28] [29] [30] with their own limitations. The approaches proposed in [27] [28] [29] only used the hand-designed features of compounds, but did not take the feature information of targets into account. Also, they generally failed to predict potential interacting compounds for a given new target (i.e., with no known interacting compound in training data), which is usually a more urgent case than predicting new compounds for targets that already have known interacting compound partners. Although a new approach has been presented in [30] to overcome these limitations, it can only be used to predict the interacting drug partners for those targets with known structures, which is not often the case.

Inspired by recent progress in representation learning and deep learning, we propose a new framework that combines unsupervised representation learning with the current powerful deep learning techniques for structure-free drug-target interaction prediction. Our method first uses latent semantic analysis [31] and Word2vec [19] [32] [20] techniques to learn the embeddings (i.e., low-dimensional representations of features) of drugs (compounds) and targets (proteins) in unsupervised manners from large compound and protein corpora, respectively. Then, given a compound-protein pair, a combination of both embeddings of compound and protein are fed into a deep neural network classifier to predict their interaction probability. We have tested our method on several benchmark datasets, including the large-scale compound-protein affinity databases (e.g., ChEMBL and BindingDB) as well as the known drug-target interactions from DrugBank. The comparisons to several baseline methods have demonstrated the superior performance of our approach in predicting new compound-protein interactions, especially when the interaction knowledge of compounds and proteins is unknown.

## 2 Methods

### 2.1 Overview

Our method consists of two main steps (Figure A1 in Appendix): (1) representation learning for both compounds and proteins; and (2) predicting interactions of compound-protein (or drug-target) pairs by a deep neural network. More specifically, in the first step, we use several natural language processing (NLP) techniques to extract useful features of compounds and proteins from the corresponding large corpora. Compounds and their basic substructures are regarded as “documents” and “words”, respectively. All possible three non-overlapping amino acid residues from a protein sequence are treated as “sentences” and “words”. Then, the embedding techniques like latent semantic analysis [31] and Word2vec [19] [32] are used to automatically learn implicit but expressive low-dimensional feature vectors of compounds and proteins from the corresponding large-scale unlabelled corpora. In the second step, a given compound-protein pair is represented by the concatenation of their embeddings of features and then fed a into deep neural network classifier to predict the likelihood of their interaction. The remaining part of this section will provide the details of individual modules, including the constructions of compound and protein features, and the deep learning network model. Implementation details are provided in Appendix (Section A2).

### 2.2 Compound feature construction

To learn good embeddings (i.e., features) of compounds, we mainly use the latent semantic analysis (a.k.a. latent semantic indexing) technique [31], which is probably one of the most effective methods for document similarity analysis [33] in the NLP field. In latent semantic analysis, each document is represented by a vector storing the term frequency or tf-idf (term frequency-inverse document frequency), which is a popularly used numerical statistic in information retrieval to describe the importance of a word in a document. Then a corpus (i.e., a collection of documents) can be represented by a matrix where each column stores the occurrences tf-idf information of terms in a document. After that, singular value decomposition (SVD) is applied to obtain a low-dimensional representation of features in documents.

In the context of compound feature extraction (Figure 1), a compound and its substructures can be viewed as a document and terms, respectively. Given a compound set *n*, we use the Morgan fingerprints [34] with radius one to scan every atom of each compound in *n* and then generate corresponding substructures. Let D denote the set of substructures generated from all the compounds in *n*. We then use a matrix *M* ϵ ℝ^|*D*|×|*N*|^ to store the word count information for these compounds, where *M*_*ij*_ represents the tf–idf of the *i*th substructure in the *j*th compound. More specifically, *M*_*ij*_ is defined as

**Figure 1:**
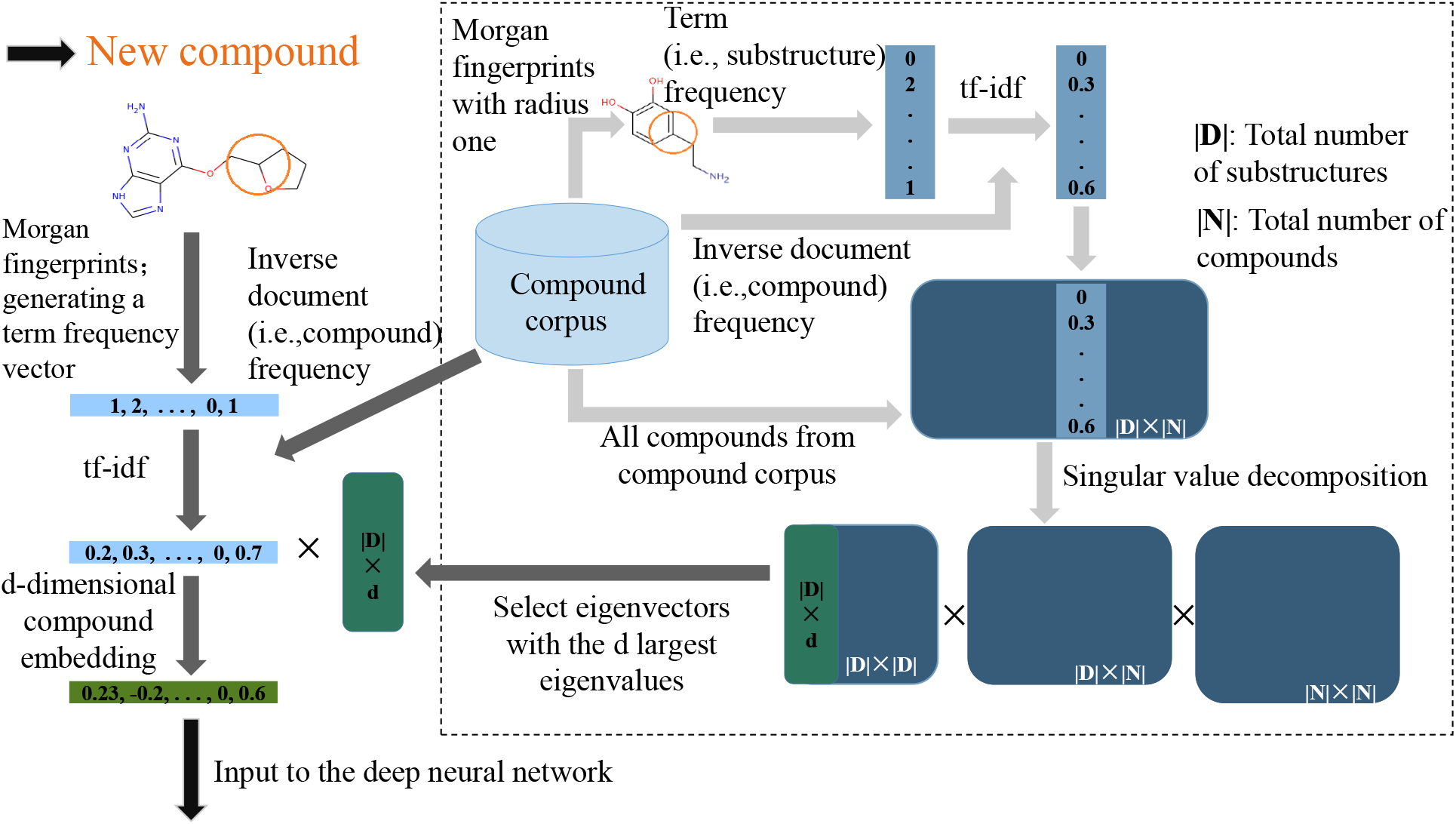
The schematic illustration of the module of compound embedding (i.e., extraction of compound features). The substructures of each compound from a compound corpus are generated by Morgan fingerprints with radius one. The term (i.e., substructure) frequency-inverse document (i.e., compound) frequency (tf-idf) information of all compounds in the corpus then forms a |*D*| × |*N*| matrix, where |*D*| and |*N*| stands for the total numbers of substructures and compounds in the corpus, respectively. Then, singular value decomposition is used to decompose this |*D*| × |*N*| matrix, and all eigenvectors corresponding to the *d* largest eigenvalues are chosen to obtain a |*D*| × *d* matrix *Û*, where *d* = 200. The matrix *Û* can be precomputed and is fixed after the above construction procedure. Then, for any new compound, its embedding is obtained from its tf-idf left multiplying by *Û*^T^. More details can be found in the main text.

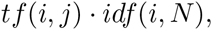

where *tf* (*i, j*) stands for the number of occurrences of the *i*th substructure in the *j*th compound, and 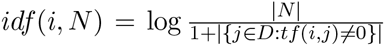. Here, |{*j* ϵ *D*: *tf*(*i, j*) ≠ 0}| represents the number of documents containing the *i*th substructure. Basically, *idf* (*i, N*) reweighs *tf*(*i, j*) so that the more common substructures will have smaller weights while those less common substructures will have larger weights. This is consistent with an observation in information theory that rarer events generally have higher entropy and thus are more informative.

After constructing matrix *M*, SVD is used to decompose *M* into three matrices *U*, Σ, *V*\* which satisfy *M* = *U*Σ*V*\* (Figure 1). Here, Σ is a |*D*| × |*N*| diagonal matrix with the eigenvalues of *M* on the diagonal, and *U* is a |*D*| × |*D*| matrix where each column *U*_*i*_ is an eigenvector of *M* corresponding to the *i*th eigenvalue Σ_*ii*_.

In order to embed compounds into a low-dimensional space ℝ^*d*^, where *d* < |*D*|, we choose the first d columns of *U* which correspond to the largest eigenvalues in Σ. Let *Û* denote the matrix with columns corresponding to the aboven chosen eigenvectors. Then the low-dimensional embeddings of *M* can be obtained by

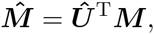

where 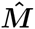 is a *d* × |*N*| matrix and its *i*th column corresponds to a *d*-dimensional embedding (or feature vector) of the ith compound. *Û*^T^ can be precomputed and fixed after being trained from a compound corpus. Given any new compound, its embedding can be obtained by left multiplying its tf-idf by *Û*^T^.

Our compound embedding framework uses the compounds retrieved from the following databases, including, all compounds labelled as active in bioassays from PubChem [16], all FDA-approved drugs from DrugBank [35], and all compounds stored in ChEMBL [36]. Duplicate compounds are removed according to their InChIs (International Chemical Identifiers). The total number of final compounds used in the compound feature extraction module to construct matrix *M* is 1,795,801. The total number of different substructures generated from the Morgan fingerprints with radius one is 18,868. We set *d* = 200, which is a recommended choice in latent semantic analysis [37].

### 2.3 Protein feature construction

To learn useful features of proteins, we consider Word2vec, a successful word embedding technique for various NLP tasks [19] [32] to embed protein sequences. In particular, we mainly use the Skip-gram with negative sampling method [32] to train the word embedding model and learn the context relations between words in sentences. We first introduce some necessary notation. Suppose that we are given a set of sentences *S* and a context window of size *b*. Given a sentence *s* ϵ *S* that is represented by a sequence of words (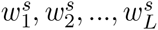), where *L* is the length of *s*, then the contexts of a word 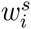 are defined as 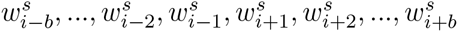. That is, all words appearing with in the context window of size *b* and centered at word *w* in a sentence are regarded as the contexts of *w*. We further use *W* to denote the set of words appearing in *S*, and #(*w, c*) to denote the total number of occurrences of word *c* ϵ *W* appearing in the *b*-sized context window of word *w* ϵ *W* in *S*. Then the Skip-gram method equips every word *w* ϵ *W* with two vectors *e*_*w*_, *a*_*w*_ ϵ ℝ^*d*^, where *e*_*w*_ is a *d*-dimensional embedding vector that we want to learn and *a*_*w*_ is a *d*-dimensional auxiliary vector. Then our goal is to maximize the following objective function:

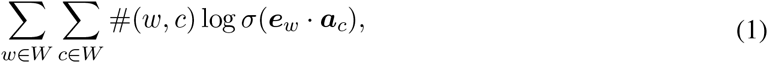

where 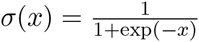 is the sigmoid function. Since the range of the sigmoid function is [0, 1], *σ*(*e*_*w*_ • *a*_*c*_) can be interpreted as the probability of word *c* being a context of word *w*, and Equation 1 can be viewed as the log-likelihood of a given sentence set *S*.

One problem in the above objective function (i.e., Equation 1) is that it does not take any negative example into account. If we arbitrarily assign any large positive values to *e*_*w*_, *a*_*c*_, *σ*(*e*_*w*_ • *a*_*c*_) would be always predicted as 1. In this case, although Equation 1 is maximized, such embeddings are surely useless. To tackle this problem, a Skip-gram model with negative sampling [32] has been proposed, in which “negative examples” *c*_*ns*_ (*c*_*ns*_ ϵ *W* and *c*_*ns*_ ≠ *w*) are drawn from a data distribution 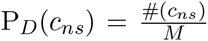, where *M* represents the total number of words in |*S* | and #(*c*_*ns*_) represents the total number of occurrences of word *c*_*ns*_ appearing in *S*. Then, the new objective function can be written as

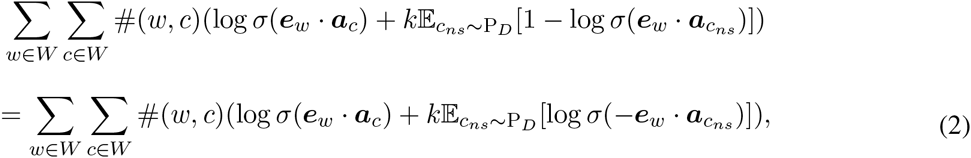

where *k* is the number of “negative examples” to be sampled for each observed word-context pair (*w, c*) during training. Maximizing this objective function can be performed using the stochastic gradient descent technique.

For each observed word-context pair (*w, c*), Equation 2 aims to maximize its log-likelihood while minimizing the log-likelihood of *k* random pairs (*w, c*_*ns*_) under the assumption that such random selections can well reflect the unobserved word-context pairs (i.e., negative examples) representing the background noise. In other words, the goal of the task is to distinguish the observed word-context pairs from the background noise distribution.

As in other existing schemes for encoding features of genomic sequences [20], each protein in our framework is regarded as a “sentence” reading from its N-terminus to C-terminus and every three non-overlapping amino acid residues can be viewed as a “word” (Figure 2). For each protein sequence, we start from the first, second and third amino acid residues from the N-terminus end, respectively, and then consider all possible “words” while discarding residues that cannot form a “word”.

**Figure 2:**
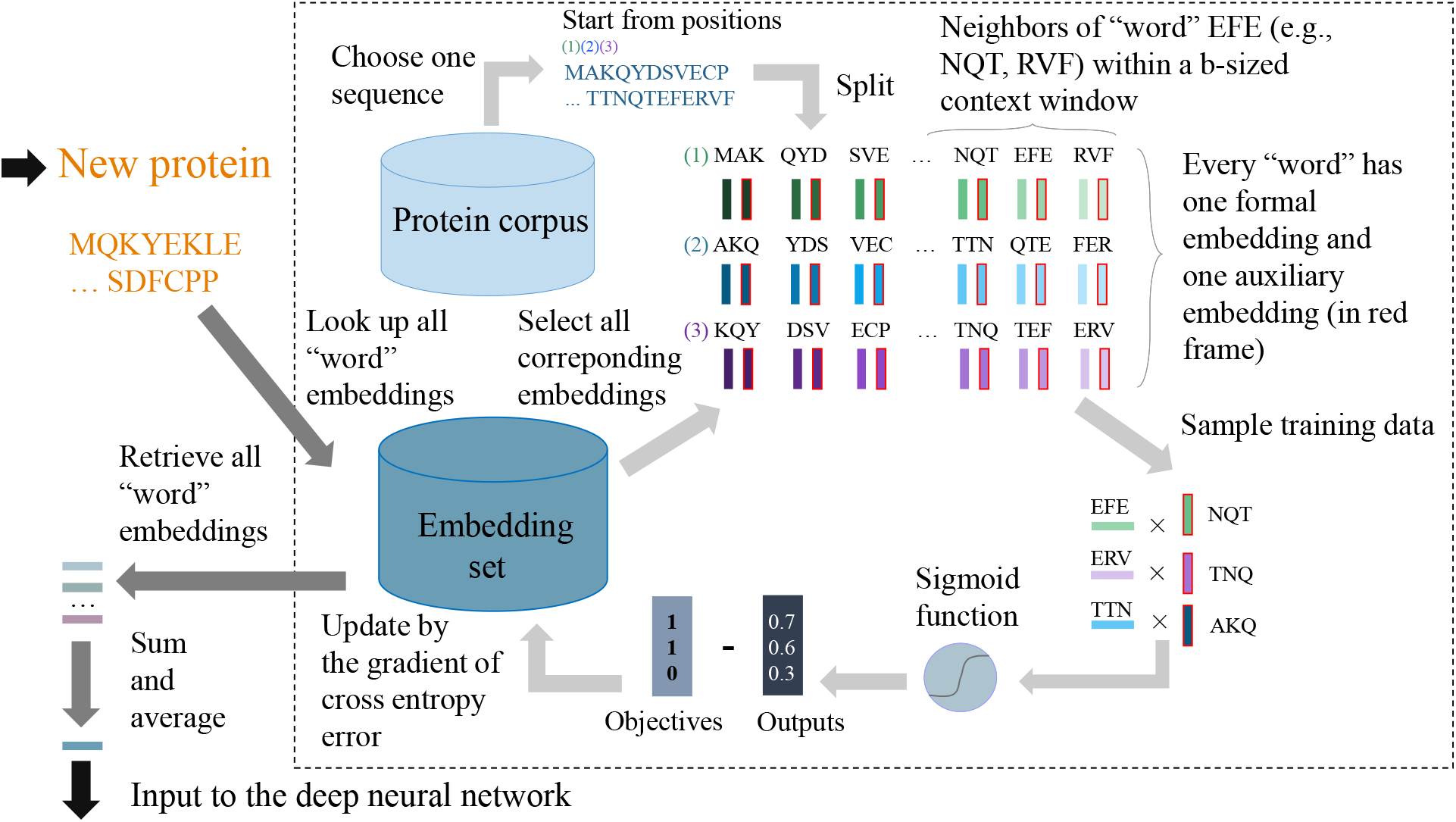
The schematic illustration of the module of protein embedding (i.e., extraction of protein features). We first build a “word” embedding set. By spliting each protein from a corpus and representing it with three “sentences”, starting from the first, second and third positions, respectively. Every three non-overlapping residues in a “sentence” is regarded as a “word”. For each “word” *w*, all “words” within a context window of size *b* and centered at *w* in any “senctence” are called the contexts (i.e., neighbors) of *w*. Every “word” is equipped with a formal embedding and an auxiliary embedding. The embeddings of “words” are learnt by maximizing the likelihood of their context words while minimizing the likelihood of random “words” given a center word. For a given protein sequence, its embedding is constructed by summing and averaging the embeddings of all “words” in its three possible encoded “sentences”. The embeddings of “words” can be precomputed and are fixed after training. More details can be found in the main text.

After converting protein sequences to “sentences” and all three non-overlapping amino acid residues to “words”. Skip-gram with negative sampling is then used to learn the embeddings of these “words”. After that, the learnt embeddings of “words” are fixed, and an embedding of any protein sequence is obtained by summing and averaging all embeddings of “words” in its all three possible encoded “sentences” (Figure 2). Note that a similar approach has been successfully used to construct text feature for text classification using Word2vec [38].

In our work, protein sequences that are used to learn the embeddings of protein features are retrieved from several databases, including PubChem [16], DrugBank [35], ChEMBL [36], Protein Data Bank [39] [40] [41] and UniProt [42]. All duplicate sequences are removed and the final number of sequences for learning embeddings is 464, 122. We follow the same principles as in [20] to choose the hyperparameters of Skip-gram. That is, the embedding dimension is set to *d* = 100, the context window size is set to *b* = 12, and the number of negative examples is set to *k* = 15, which was also used in [32].

### 2.4 The deep neural network model

Suppose that we are given a training data set {(*c*_*i*_, *p*_*i*_) | *i* = 1, 2, …, *n*} and a corresponding label set { *y*_*i*_ | *i* = 1, 2, …, *n*}, where *N* stands for the total number of compound-protein pairs, *y*_*i*_ = 1 means that compound *c*_*i*_ and protein *p*_*i*_ bind to each other, and *y*_*i*_ = 0 otherwise. We first use the schemes described in Sections 2.2 and 2.3 to construct features for individual compounds and proteins. For every compound-protein pair (*c*_*i*_, *p*_*i*_), *c*_*i*_ and *p*_*i*_ are represented by a 200-dimensional embedding and a 100-dimensional embedding, respectively. We then concatenate these two embeddings into a 300-dimensional vector as the final input to our deep learning model (i.e., deep neural network).

Our deep neural network (DNN) is a sequence of fully-connected layers that take the concatenated 300-dimensional feature vector of each compound-protein pair as input and classify this pair as binding or non-binding. The basic DNN architecture consists of an input layer *L*_0_, an output layer *L*_*out*_ and *H* hidden layers *L*_*h*_ (*h* ϵ {1, 2, …, *H*}) between input and output layers. Each hidden layer *L*_*h*_ is a set of several units which can be arranged as a vector 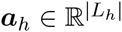, where |*L*_*h*_| stands for the number of units in *L*_*h*_. Then, each hidden layer *L*_*h*_ can be parametrized by a weight matrix 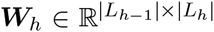, a bias vector 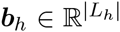 and an activation function *f* (•). More specifically, the units in *L*_*h*_ can be calculated by

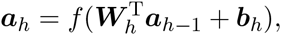

where *h* = 1, 2, …, *H*, and the units *a*_0_ in the input layer *L*_0_ are from the 300-dimensional feature vector of a compound-protein pair. The activation function is chosen as *f* (*x*) = max(0, *x*), which is also known as the Rectified linear unit (ReLU) fucntion [43]. ReLU is a common choice of activation function in deep learning [44], probably due to its sparsity property, high computational efficiency and no gradient vanishing effect during back-propagation training [45].

After we calculate *a*_*H*_ for the final hidden layer, the output layer *L*_*out*_ is simply a logistic regression model that takes *a*_*H*_ as its input and computes

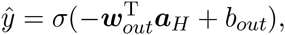

where the output 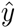 is the confidence score of the predicted binding between the given compound-protein pair, *σ* is the sigmoid function, 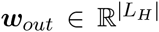 and *b*_*out*_ ϵ ℝ are parameters of *L*_*out*_. Since the sigmoid function has a range [0, 1], 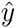 can be interpreted as the binding probability of the given compound-protein pair.

To learn *w*_*out*_, *b*_*out*_ and all parameters *W*_*h*_, *b*_*h*_ in hidden layers from the training data set and the corre-sponding label set, we want to minimize the following cross entropy loss:

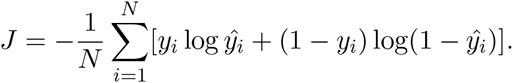

The above minimization problem can be solved using the stochastic gradient descent and back-propogation techniques. In addition, we apply two popular tricks that have been widely used in the deep learning comm-nuity, including dropout [46] and batch normalization [47], to further enhance the performance of our DNN. Dropout sets units in hidden layers to zero with probabiltiy *p*. This technique can effectively alleviate the potential overfitting problem in deep learning [22] [46]. Batch normalization normalizes the outputs of units in a hidden layer to zero mean and unit standard deviation. Such an operation can accelerate the training process and act as a regularizer [47].

Since the positive examples and negative examples are possibly imbalanced, our classifier may learn a “lazy” solution. That is, in such a case of skewed data distribution, the classifier would easily predict the dominant class given any input. To avoid this problem, we resample the examples from the minor class so that the number of positive examples and the number of negative examples are comparable to each other during training. In our computational tests, we also implement an ensemble version of DNN and use the average prediction over ten DNNs to obtain relatively more stable prediction results.

## 3 Results

### 3.1 Performance evaluation on compound-protein interactions

Databases like ChEMBL [36], BindingDB [48] and PDBbind-CN [49] [50] [51] have collected a large amount of binding profiles (e.g., binding affinities or potencies) between compounds and proteins. We first evaluated our method on the compound-protein pairs extracted from these databases. In particular, we mainly used the bioactivity data retrieved from ChEMBL [36] to evaluate our model. Compound-protein pairs with half maximal inhibitory concentration (IC50) or inhibition constant (Ki) ≤ 1*µM* were chosen as positive examples, while pairs with IC50 or Ki ≥ 30*µM* were chosen as negative examples. This resulted in 360,835 positive examples and 93,903 negative examples.

To justify our criterion of choosing positive and negative examples, we mapped the known interacting drug-target pairs extracted from DrugBank [35] (retrieved in November 11, 2015) to the corresponding compound-protein pairs in ChEMBL. A drug-target pair from DrugBank was said to be in ChEMBL if there exist compound-target pairs in ChEMBL that have identical InChIs for compounds or drugs and sequence identity score *t* ≥ 0.40 for proteins. Given two protein sequences *s* and *s*′, sequence identity score *t* is defined as 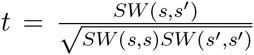 [13], where *SW*(•, •) is the Smith-Waterman score [52]. As shown in Figure A2 (in Appendix), the binding affinities or potencies (measured by Ki or IC50, respectively) of most drug-target pairs lied in the range of [0*µM*, 1*µM*]. Also, 1*µM* was used as an good indicator for identifying strong binding affinities in the literature, e.g., [53]. Therefore, here we considered ≤ 1*µM* of IC50 or Ki as a reasonable criterion for choosing positive examples. Since there is no well-defined dichotomy between high and low binding affinities, we used a threshold ≥ 30*µM* that is relatively far away from 1*µM* to choose negative examples. Note that the same choice was also used in [30] to choose negative examples.

As pointed out in [54], cross-validation may not be a proper way to evaluate prediction performance if training dataset contains many compounds or proteins with only one interaction, especially for leave-one-out cross-validation (LOOCV). In such a case, training methods may lean to exploit the bias towards those proteins or compounds with single interactions to boost the performance of LOOCV. As suggested in [54], choosing a small *k* for *k*-fold cross-validation instead of using LOOCV can mitigate this effect. The unique interactions were also suggested to separate from the non-unique interactions during the validation process [54]. Given a compound-protein pair from a dataset, if the compound or protein only appeared in this pair, we considered this pair as unique. Among 360,835 positive examples and 93,903 negative examples obtained from ChEMBL, there were 156,083 unique positive examples and 39,857 unique negative examples. During our validation tests, we also used non-unique examples as training data and tested the prediction performance on unique examples, and vice versa.

We further defined a compound-protein pair as double unique if both the protein and the compound appeared only in this pair. This resulted in 78 double unique examples labelled as positive and 59 double unique examples labelled as negative. In an additional validation test, all non-unique examples were used as training data and then the trained model was evaluated on these double unique examples. Such a scenario was equivalent to predicting compound-protein pairs in which the previous interaction knowledge of the involving compound and protein is entirely unknown.

In all computational experiments, random forest [55], one of the most powerful classifiers [56], was also used as a baseline method to demonstrate the need to use the deep neural network model. Although our deep neural network only slightly outperformed random forest in the first two test settings (Figures 3a-3b), we observed significant AUC improvement from our approach (0.7890 versus 0.5985) in the third test setting (Figure 3c). This result suggested that our deep learning model were much more capable of predicting new compound-protein pairs whose interaction knowledge is entirely unknown. In general, the prediction task in this setting is much more challenging than the other situations, in which the interaction knowledge of the involving compound or protein is available. On the other hand, our approach still achieved decent prediction performance (AUC of 0.7890) in this tough situation, which demonstrated that our model can still preserve generalization ability and predictive power for predicting novel compound-protein pairs when their previous interaction knowledge is completely unknown.

In addition to ChEMBL, we performed two additional tests and further assessed the performance of our method on two external datasets: BindingDB [48], which stores the binding affinities between proteins and drug-like small molecules, and PDBbind-CN [49], which collects the experimentally measured binding affinity data from the Protein Data Bank [39] [40] [41]. When evaluating on the BindingDB dataset, we used the same criterion (i.e., IC50 or Ki ≤ 1*µM* for positive examples and ≥ 30*µM* for negative examples) to label compound-protein pairs, which resulted in 418,577 and 117,210 positive and negative examples, respectively. In the additional tests, we used compound-protein pairs from ChEMBL as training data, while those from BindingDB as test data. Also the overlaps between BindingDB and ChEMBL were all removed from training data. Here, if a compound-protein pair from ChEMBL had identical compound InChIs and protein sequence identity score *t* ≥ 0.40 with pairs from BindingDB, it was regarded as a overlap and removed from training data. This resulted in 75,740 positive examples and 26,785 negative examples in the refined training dataset. We also tested the situation in which overlaps were removed from BindingDB instead of ChEMBL. This resulted in 128,415 positive examples and 50,100 negative examples in the refined test dataset.

We also evaluated our prediction method on PDBbind-CN, which has about 25 % overlaps with Bind-ingDB. After removing overlaps from ChEMBL, we had 359,743 positive examples, 93,737 negative examples in the training data and 2,188 positive examples and 578 negative examples in the test data.

The evaluation results on the BindingDB and PDBbind-CN datasets are summarized in the second row of Figure 3. Again, our deep learning model outperformed random forest in most cases, especially on PDBbind-CN (with 0.8796 versus 0.7626 on AUC, and 0.8030 versus 0.7863 on ACC). In addition, the performance evaluated on BindingDB and PDNbind-CN agreed well with the test results on ChEMBL itself, which also indicated a good generalization ability of our method.

**Figure 3:**
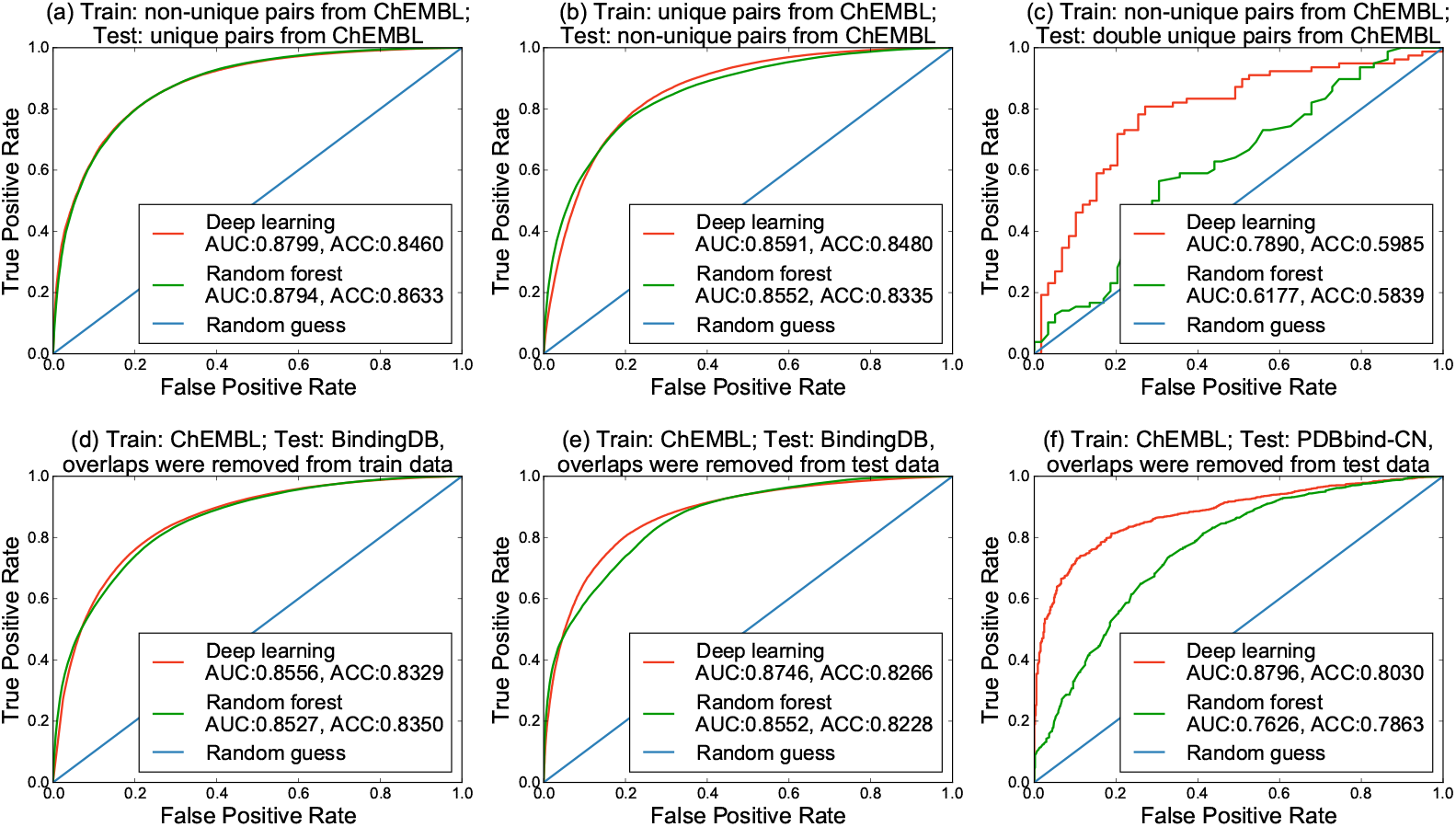
Performance evaluation on the compound-protein pairs from ChEMBL, BindingDB and PDBbind-CN. All embeddings are precomputed as described in Section 2.

As summarized in Figure 3, all the above tests suggested that our method surpassed random forest and can generalize well even when the compound and protein from a to-be-predict pair are completely not seen in training data. To see how the deep neural network extracted high-level abstractions of features from input data, we used t-SNE [57] to visualize and compare the distributions of positive and negative examples on their 300-dimensional input features and the features represented by the last hidden layer in the deep neural network. Here, the deep neural network was trained on ChEMBL and 5,000 positive and 5,000 negative examples randomly selected from BindingDB were serviced as the test data and used for the visualization. The visualization (Figure 4) showed that the data were better organized by the deep neural network so that the final output layer (which was simply a logistic regression classifier) can better exploit the hidden features to yield improved classification results.

**Figure 4:**
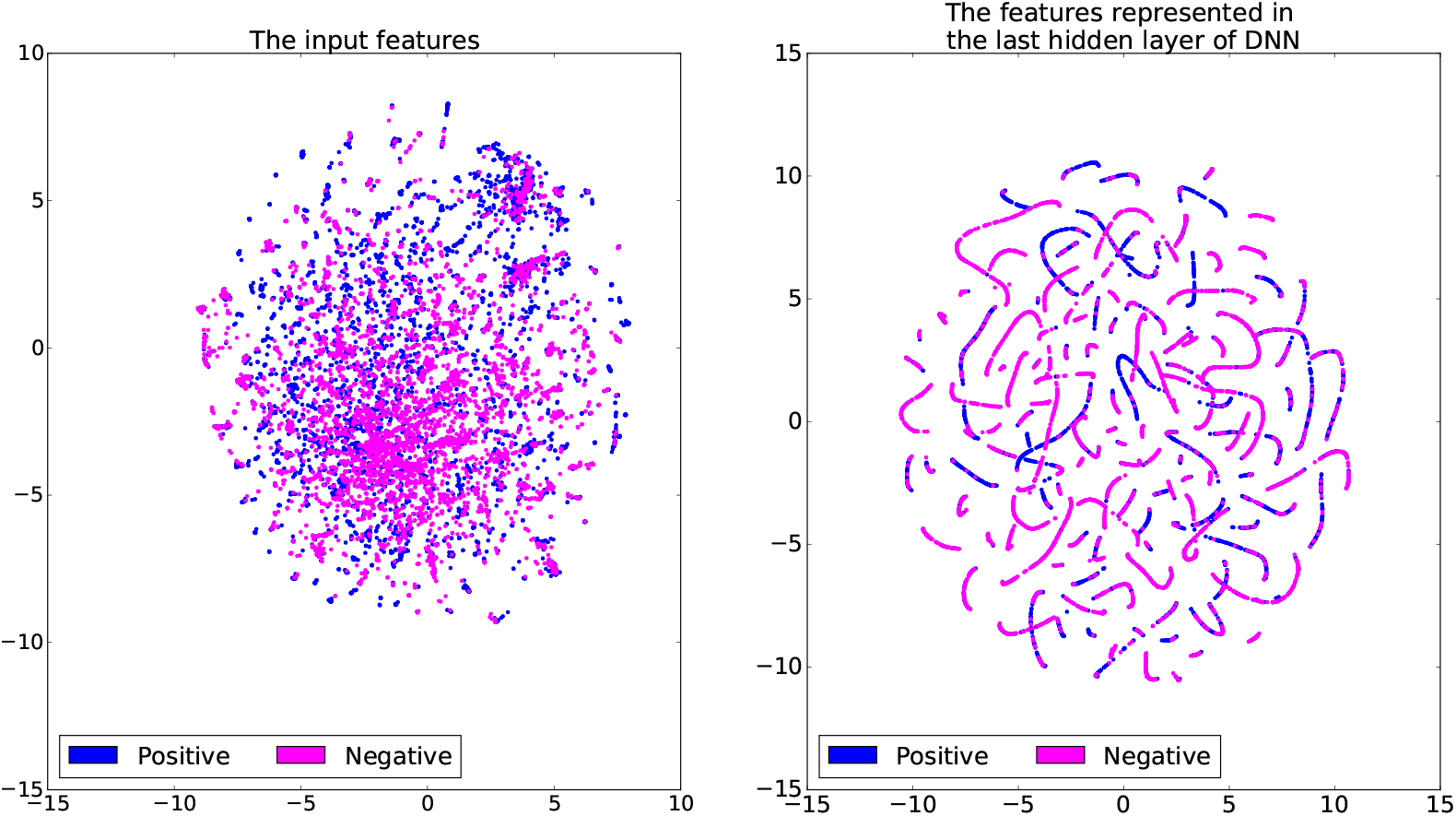
Visualization of the distributions of positive and negative examples by t-SNE [57] on the 300-dimensional input features of DNN (left) and the feautres represented by the last hidden layer of the DNN (right). DNN was trained on the ChEMBL dataset. 5,000 positive and 5,000 negative examples were randomly selected from BindingDB and serviced as the test data, which were used in the visualization.

We also compared our method to AtomNet [30] (a structure-based deep learning approach for compound-protein binding potency prediction) on DUD-E [58], a benchmark dataset for evaluating the molecular docking programs. The tests on the DUD-E dataset also showed that our approach had superior prediction performance over existing methods in various settings (see Section A1 from Appendix for more details).

### 3.2 Performance evaluation on drug-target interactions

Next, we tested our method on the known drug-target interactions obtained from DrugBank [35]. Here, we regarded all drugs and targets that were stored in DrugBank before November 11, 2015 as “old” drugs and “old” targets respectively, while those drugs and targets stored after November 11, 2015 were considered as “new” drugs and “new” targets, respectively. We then defined four datasets, including Data-1, Data-2, Data-3 and Data-4, which corresponded to all interactions between old drugs and old targets, between new drugs and old targets, between old drugs and new targets, and between new drugs and new targets, respectively. Basic statistics of these four datasets can be found in Table A2 in Appendix.

We then used Data-1 as training data for our method, and evaluated its performance based on Data-2, Data-3 and Data-4, respectively. Here, the drug-target pairs that were considered as positive examples, if they have interaction records in the given dataset, and were defined as negative examples otherwise. Here, the training examples were heavily imbalanced (i.e., 9, 144 positive examples versus 1,390 × 1,606 − 9,144 = 2,207,136 negative examples). We applied the resampling scheme as described in Section 2.4 to make sure that the numbers of positive and negative examples were roughly the same. The test results showed that our deep neural network achieved much more robust performance than random forest (Figure 5). Although both deep learning and random forest methods achieved similar evaluation results on Data-2 (i.e., interactions between “new” drugs and “old” targets), the deep neural network model yielded much better prediction results than random forest when evaluating on Data-3 (i.e., interactions between “old” drugs and “new” targets) and Data-4 (i.e., interactions between “new” drugs and “new” targets) (Figures 5b-5c). Together with the previous results presented in Section 3.1, these test results implied that although both approaches were equipped with the extracted features obtained from the embedding techniques (as described in Section 2), our deep learning method had better generalization capacity than random forest to achieve more robust prediction results, especially for new drug-target pairs with partial or no interaction information. Compared to the previous results on predicting compound-protein pairs without known interaction knowledge (Figure 3c), the prediction results on the interactions between “new” drugs and “new” targets here were much worse. This was probably due to the much smaller size and much more imbalanced distribution of the drug-target interaction training data obtained from DrugBank.

**Figure 5:**
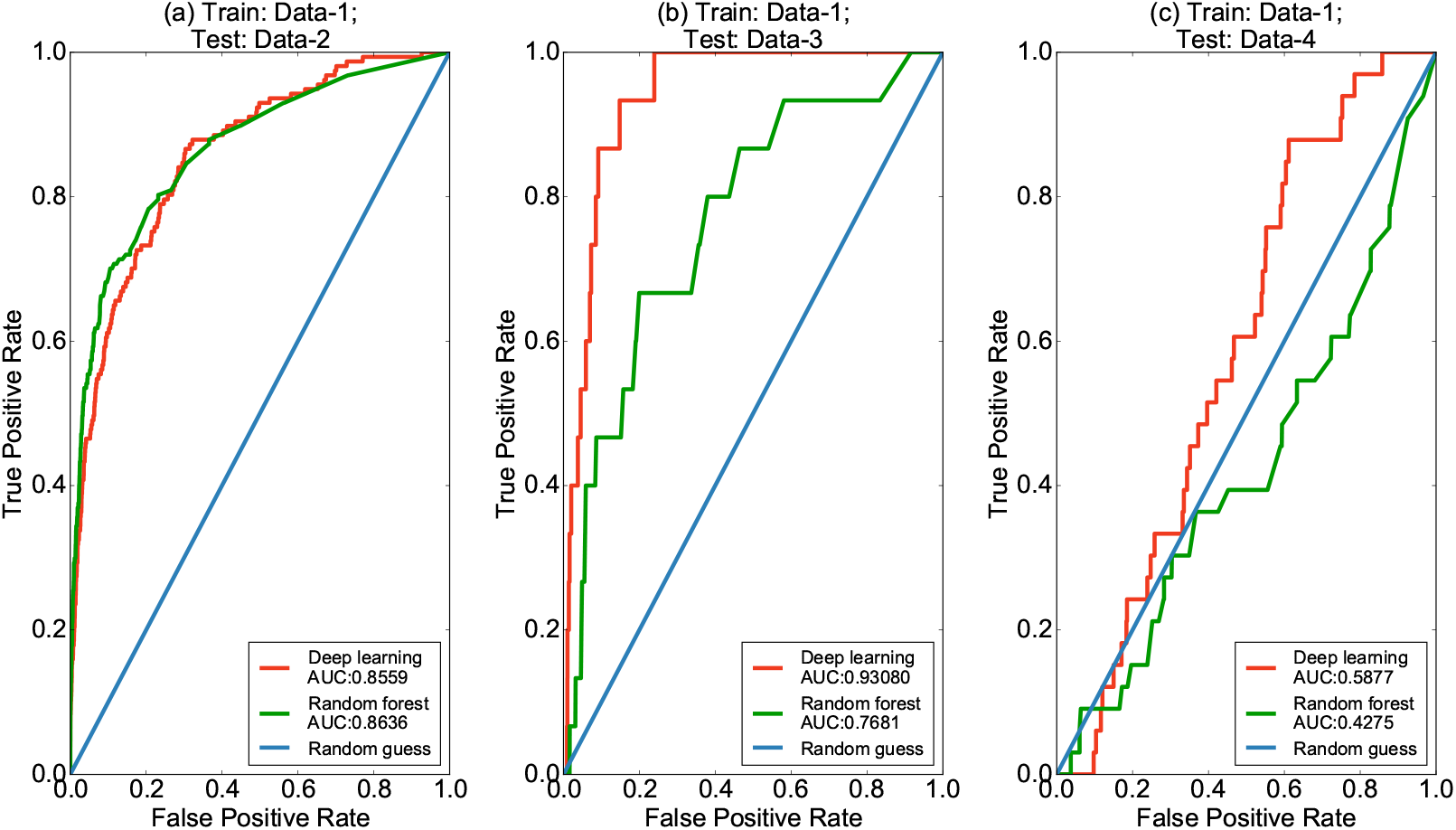
Test results on the known drug-protein interactions obtained from DrugBank by the deep neural network and random forest. In all three tests, we used Data-1 (i.e., interactions between “old” drugs and “old” targets) as training data, and evaluated the prediction methods based on Data-2 (i.e., interactions between “new” drugs and “old” targets), Data-3 (i.e., interactions between “old” drugs and “new” targets) and Data-4 (i.e., interactions between “new” drugs and “new” targets), which correspond to (a), (b), (c), respectively. All embedding features were precomputed, as described in Section 2.

### 3.3 Novel drug-target interaction predictions

We also used all known interaction data from DrugBank [35] to train our deep learning model and then examined the novel prediction results on those missing interactions, i.e., the drug-target pairs that did not have any known interaction record in DrugBank. We particularly looked into the list of top predictions with the highest rating scores, and checked whether we can find the known experimental studies in the literature to support these predictions. Note that, here, we excluded those “easy” predictions, in which the targets were from the same protein family or had sequence identity score ≥ 0.40 from any protein in the training data.

We that most of the top predictions with the highest prediction scores can be supported by the known evidence in the literature. For example, among the top 10 predictions, six novel drug-protein interactions were consistent with the previous studies in the literature (Table A3 in Appendix). Here we describe several examples of these novel predictions. First, DrugBank [35] showed that norethisterone, a drug often used for hormonal contraception, only acts on the progesterone receptor. Our new predictions indicated that norethisterone can also interact with the estrogen receptor (ER), an important nuclear receptor binding to the sex hormone estrogen. This prediction was consistent with the previous known result that norethisterone can bind to ER and induce the ER-activating functions [59] [60]. Also, our method predicted that cefamandole (which is a cephalosporin antibiotic) can interact with serum albumin. This prediction can be supported by a previous work on studying the binding characteristics of cefamandole with bovine serum albumin [61] and another known observation that cefamandole can help the clearance of thiopental from human serum albumin [62]. In addition, our model predicted that amitriptyline, a drug commonly used to treat many mental diseases, such as major depressive disorder and anxiety disorder, can have an interaction with the dopamine receptor. This result can be supported by a previous known evidence that amitriptyline upregulates both DRD2 (i.e., dopamine receptor D2 isoform) and DRD3 (i.e., dopamine receptor D3 isoform) [63]. Moreover, according to our prediction results, trilostane, an inhibitor of 3*β*-hydroxysteroid dehydrogenase for treating the Cushing's syndrome, can bind to the androgen receptor, an important therapeutic target in prostate cancer. Such a predicted interaction agreed with a previous experimental observation [64] that trilostane can induce agonistic activity on the androgen receptor in human prostate cancer cells. Our predictions also showed that benzphetamine can act on amine oxidases, which can be supported by the known facts that benzphetamine is a substituted amphetamine and amphetamine can be used an effective inhibitor of copper-containing amine oxidases [65].

## 4 Discussion

In this paper, we propose a new and scalable framework that combines data-driven representation learning with deep learning to predict new drug-target (or compound-protein) interactions. By leveraging large-scale unlabelled data of compounds and proteins, our embedding schemes extract low-dimensional expressive features from raw data without the requirements of available protein structure and any known interaction information. The combination of effective feature embedding strategies and powerful deep learning models is particularly useful for fully exploiting the massive amount of compound-protein binding data available from large-scale databases, such as PubChem and ChEMBL. The effectiveness of our method has been validated on several large-scale compound-protein binding datasets as well as the known interactions between FDA approved drugs and targets.

Recently we have been collaborating with the National Center for Drug Screening, the Chinese Academy of Sciences, on experimentally validating a number of our predicted interactions between several G proteincoupled receptors (GPCRs) and small molecule compounds. The preliminary validation results have shown that more than ten compounds among hundreds of predictions can have significant inhibitory effects on the GLP-1 receptor (GLP-1R), according to the measured cAMP response. GLP-1R belongs to the glucagon subfamily of class B GPCRs, and is an important target involving type 2 diabetes. Thus, identification of the effective drug-like compounds acting on GLP-1R will be extremely useful in drug development for the treatment of type 2 diabetes. Currently our collaborators are working on conducting more wet-lab experiments, such as functional assays and receptor binding assays, to further validate these prediction results.

## Appendix

## A1 Performance evaluation on the DUD-E benchmark dataset

We also compared our method to AtomNet [30] (a structure-based deep learning approach for compound-protein binding potency prediction) on the DUD-E [58], which is a benchmark dataset widely used for evaluating the molecular docking programs and contains 22,886 active compounds against 102 targets. Each active compound in DUD-E is paired with 50 decoys that share similar physico-chemical properties but with dissimilar 2D topologies under the assumption that such dissimilarity in compound structure will result in different pharmacological activities with high probability.

We adopted two experiment settings from [30] to make comparisons. More specifically, in the first setting, cross-validation was done for 102 proteins, i.e., the data were separated according to proteins. In the second setting, cross-validation was done for all pairs, i.e., all compound-protein pairs were divided into three groups for validation. In addition to Atomnet, we compared our method to random forest.

AUC and ACC under two different settings among three methods were shown in Table S1. Our deep learning model outperformed both random forest and AtomNet on both settings. More specifically, our structure-free embedding features with deep learning model outperformed AtomNet which required protein structures and adopted a convolutional neural network for classification with large margin, which demonstrated the superiority of our embedding features. In addition, it is noticeable that our deep learning model achieved significantly better AUC than random forest in the first experiment setting. Since the first setting was generally harder than the second one as protein information was not seen by classifiers during validation, this result indicated that our deep learning model may have better generalization ability than random forest.

## A2 Implementation

The Morgan fingerprints of compounds are generated by RDKit [66]. Latent semantic analysis and Word2vec (Skip-gram with negative sampling) are implemented by Gensim [67], a Python library designed to automatically extract semantic topics from documents. Our DNN implementation is based on Keras [68], a highly modular deep learning library. For all experiments, we use six hidden layers with 600, 256, 128, 64, 32 and 16 units, respectively. We set the dropout rate *p* = 0.2. Batch normalization is added to all hidden layers.

**Figure A1:**
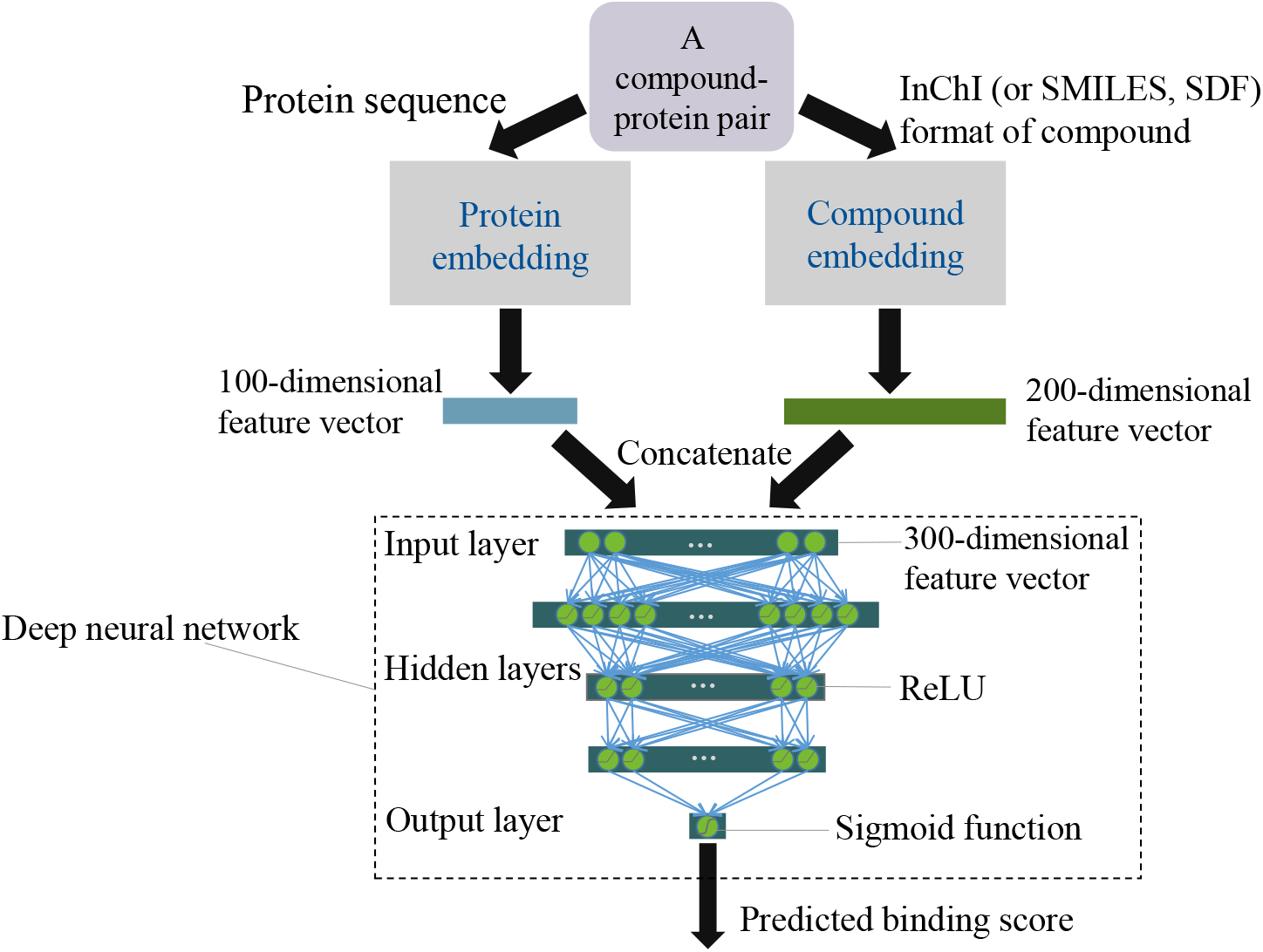
The schematic workflow of our prediction model. To predict the binding score (i.e., the probability of interaction) of a given compound-protein pair, protein sequence and the compound InChI (or SMILES and SDF) format are provided as inputs to the modules of protein and compound embeddings, respectively. A 100-dimensional vector of protein features and a 200-dimensional vector of compound features obtained from these two embedding modules are then concatenated together to form a 300-dimensional input vector to a deep neural network classifier, which predicts the likelihood of the interaction for the given compound-protein pair.

**Table A1:**
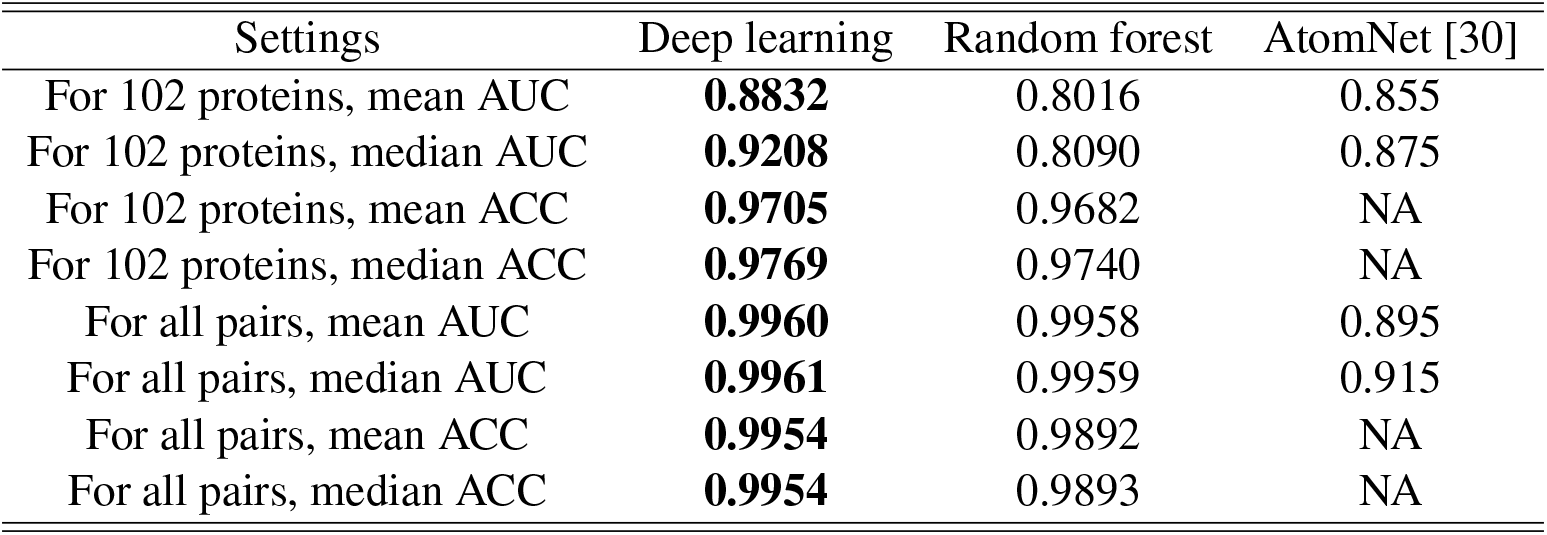
The 3-fold cross-validation (CV) reults on DUD-E. In the first setting, corresponding to the first to fourth rows, cross-validation was performed for 102 proteins, i.e., the data were separated according to proteins. In the second setting, corresponding to the fifth to eighth rows, cross-validation was conducted for all pairs, i.e., all compound-protein pairs were divided into three groups for validation. AUC and ACC were evaluated for our deep neural network as well as AtomNet [30] and random forest. All embedding features were precomputed and fixed, as described in Section 2. The best result in each test is shown in bold.

| Settings | Deep learning | Random forest | AtomNet [30] |
| --- | --- | --- | --- |
| For 102 proteins, mean AUC | **0.8832** | 0.8016 | 0.855 |
| For 102 proteins, median AUC | **0.9208** | 0.8090 | 0.875 |
| For 102 proteins, mean ACC | **0.9705** | 0.9682 | NA |
| For 102 proteins, median ACC | **0.9769** | 0.9740 | NA |
| For all pairs, mean AUC | **0.9960** | 0.9958 | 0.895 |
| For all pairs, median AUC | **0.9961** | 0.9959 | 0.915 |
| For all pairs, mean ACC | **0.9954** | 0.9892 | NA |
| For all pairs, median ACC | **0.9954** | 0.9893 | NA |

**Figure A2:**
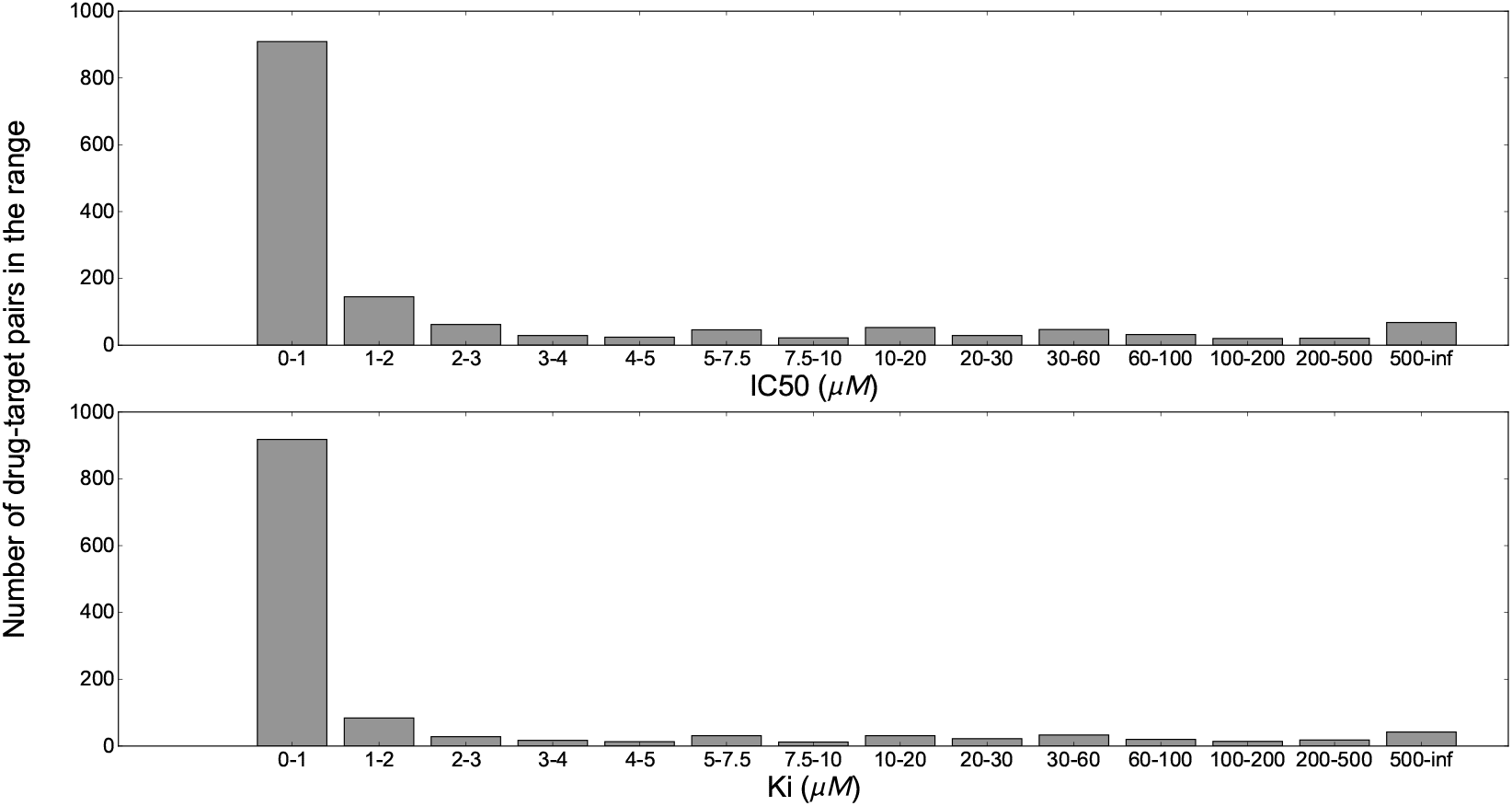
Mapping the known interacting drug-target pairs from DrugBank to the correponding compound-protein pairs in ChEMBL. We plot the ranges of the recorded values of IC50 and Ki in ChEMBL and their corresponding numbers of known interacting drug-target pairs that were mapped from DrugBank.

**Table A2:** Basic statistics of four different datasets obtained from DrugBank [35]. Drugs and targets stored in DrugBank before November 11, 2015 are regarded as “old” drugs and “old” targets, while those drugs and targets stored after November 11, 2015 are regarded as “new” drugs and “new” targets. Data-1 contains all old drugs, old targets and their known interactions. Data-2 contains all new drugs, old targets and their known interactions. Data-3 contains all old drugs, new targets and their known interactions. Data-4 contains all new drugs, new targets and their known interactions.

|  | Data-1 | Data-2 | Data-3 | Data-4 |
| --- | --- | --- | --- | --- |
| #Drugs | 1380 | 54 | 1380 | 54 |
| #Targets | 1606 | 1606 | 51 | 51 |
| #Interactions | 9144 | 157 | 15 | 33 |

**Table A3:** Six novel drug-target interaction predictions among the top 10 results with the highest prediction scores that can be supported by previous studies in the literature.

| Drug ID on DrugBank | Drug name | Protein UniProt ID | Protein name | Supporting references |
| --- | --- | --- | --- | --- |
| DB00717 | Norethisterone | P03372 | ESR1 | [59] [60] |
| DB00321 | Amitriptyline | P14416 | DRD2 | [63] |
| DB00321 | Amitriptyline | P21728 | DRD1 | [63] |
| DB01326 | Cefamandole | P02768 | ALB | [62] [61] |
| DB01108 | Trilostane | P10275 | Androgen receptor | [64] |
| DB00865 | Benzphetamine | P27338 | Amine oxidase | [65] |

